# C-terminal CDC42 variants in autoinflammatory patients specifically trigger actin defects and NF-κB hyperactivation

**DOI:** 10.1101/2024.06.26.600829

**Authors:** Alberto Iannuzzo, Philippe Mertz, Selket Delafontaine, Rachida Tacine, Guilaine Boursier, Véronique Hentgen, Sophie Georgin-Lavialle, Isabelle Meyts, Jérôme Delon

**Affiliations:** Université Paris Cité, CNRS, Inserm, Institut Cochin, F-75014 Paris, France; Laboratory for Inborn Errors of Immunity, Department of Pediatrics, KU Leuven, UZ Leuven, 3000 Leuven, Belgium; Department of Pediatrics, University Hospitals Leuven, Leuven, Belgium; Laboratoire de Génétique des Maladies Rares et Autoinflammatoires, Service de Génétique Moléculaire et Cytogénomique, National Reference Center for Autoinflammatory Diseases and AA Amyloidosis, Centre Hospitalier Universitaire Montpellier, Université de Montpellier, Montpellier, France; Department of General Pediatrics, Centre Hospitalier de Versailles, Le Chesnay, France; Department of Internal Medicine, National Reference Center for Autoinflammatory Diseases and Amyloid A Amyloidosis, Tenon Hospital, Sorbonne Université, Paris, France

**Keywords:** CDC42 GTPase, mutations, autoinflammation, actin, NF-κB

## Abstract

**Background:** CDC42 belongs to the RHO GTPases family. Recently, four variants were identified in autoinflammatory patients. One variant affects the N-terminal part of the protein while the three others are located in the C-terminal region. To date, most of the functional defects were only reported for the C-terminal R186C variant. The other three variants are far less characterized at the functional level.

**Objectives:** We aimed to investigate whether all four CDC42 variants share common signaling alterations.

**Methods:** We performed in depth imaging analysis of actin cytoskeleton and NF-κB nuclear translocation, coupled to flow cytometry in cells from patients or in the monocytic THP-1 cell line.

**Results:** We show that the N-terminal Y64C CDC42 variant localizes normally in cells and does not exhibit any defect in actin filaments formation or NF-κB activation. By contrast, all three C-terminal CDC42 variants have aberrant subcellular localizations and share common functional alterations. They exhibit a strong reduction or complete block in their abilities to polymerize actin filaments. They also show more NF-κB nuclear translocation and phosphorylation. However, we suggest that there is no causal relationship between these two events. Artificial reduction in cellular actin content using specific pharmacologic drugs is indeed not sufficient to hyperactivate NF-κB.

**Conclusions:** This study further extends the spectrum of defects observed in autoinflammatory CDC42 patients, and pinpoints a functional heterogeneity between N- and C-terminal CDC42 variants. We also show that CDC42 patients should not be necessarily classified among actinopathies. Altogether, the functional defects we report here can lead the way towards more personalized therapeutic interventions.

## INTRODUCTION

CDC42 is one of the best-characterized members of the RHO GTPases family. Its involvement in the regulation of the actin cytoskeleton and cell polarity are its most studied functions (1). In this context, CDC42 also plays a role in a wide variety of actin cytoskeleton-dependent cellular processes such as cytokinesis, phagocytosis, cell migration, morphogenesis, and chemotaxis. Membrane binding of CDC42 requires the post-translational addition of lipids to the cysteine residue located in the C-terminal CAAX sequence by geranylgeranyl transferase. This process is called prenylation (2).

Heterozygous pathogenic variants in *CDC42* have been discovered in several patients presenting complex syndromes (3). One such rare developmental disorder is the Takenouchi-Kosaki syndrome (TKS) whose hallmarks are growth and intellectual delay, structural brain abnormalities with sensorineural deafness, hypothyroidism, dysmorphisms, camptodactyly, and recurrent infections (4–6). All these initial patients with TKS share a unique Y64C mutation in the N-terminal part of CDC42 (Fig. 1A). Martinelli *et al*. have then described fourteen additional patients carrying mostly N-terminal mutations, including Y64C in CDC42, and presenting a group of clinically heterogeneous phenotypes characterized by variable growth deregulation, facial dysmorphia, neurodevelopmental defects, immunological and hematological abnormalities (7) (Table 1). More recently, variants in CDC42 have been associated with complex auto-inflammatory syndromes. This way, a TKS patient carrying a Y64C mutation was reported with late-onset systemic inflammation and myelofibrosis (8). In parallel, we and others have described a recurrent R186C mutation in the C-terminal domain of CDC42 in a larger number of patients with neonatal-onset severe autoinflammation with increased serum pro-inflammatory cytokines, hepatosplenomegaly and hemophagocytic lymphohistiocytosis (HLH), pancytopenia and rash (NOCARH syndrome) (9–15) (Fig. 1A, Table 1). It has been shown that this R186C variant mutated at the vicinity of the C-terminal CAAX sequence carries an aberrant palmitoylation in position 186 that is responsible for its trapping in the Golgi apparatus (9, 11, 14, 16). This suggests that this variant is subject to a dual lipid modification: in addition to the geranylgeranyl lipid moiety present in the Cys188 of the CAAX sequence of WT CDC42, CDC42 R186C exhibits an additional palmitoylation in position 186. Interestingly, others have reported additional patients carrying a C188Y variant in CDC42 (Fig. 1A, Table 1) with a clinical phenotype reminiscent of the R186C patients’ phenotype (10, 15). The mutation of the Cys188 of the CAAX sequence precludes the geranylgeranylation of this variant that thus carries no lipid modification and consequently, cannot bind membranes. Lastly, the mutation of the STOP codon of CDC42 that adds up a stretch of 24 amino acids has also been reported in autoinflammatory patients (10, 17). This *192C*24 variant carries two palmitoyl modifications and is also trapped in the Golgi apparatus (16). Altogether, it is puzzling that these three C-terminal CDC42 variants (Fig. 1A) classified in the same group and all identified in severe and early onset autoinflammatory syndromes patients exhibit an excess or an absence of lipid modification.

**TABLE 1.** Clinical details of autoinflammatory CDC42 patients reported to date.

|  | Y64C<br>% (data available for N/<br>total patients) | R186C<br>% (data available for N/<br>total patients) | C188Y<br>% (data available for N/<br>total patients) | 192Cys*24<br>% (data available for N/ total<br>patients) |
| --- | --- | --- | --- | --- |
| Number of described patients | 8 | 13 | 4 | 2 |
| Gender | 7 F/ 1 M | 5 F/ 8 M | 3 F/ 1 M | 1F / 1 M |
| Age at onset | Neonatal | Neonatal | Neonatal | Neonatal – Childhood |
| Delayed neurodevelopment | 100 (8/8) | 0 (0/5) | 100 (1/1) | 50 (1/2) |
| Failure to thrive | 100(7/7) | 80 (8/10) | 100 (3/3) | 100 (1/1) |
| Dysmorphic features | 100 (7/7) | 41.7 (5/12) | 75 (3/4) | 100 (2/2) |
| Recurrent infections | 50 (4/8) | 11.1 (1/9) | 66.7 (2/3) | 100 (2/2) |
| Hypogammaglobulinemia | 66.7 (4/6) | 0 (0/8) | NA | NA |
| Fever | 12.5 (1/8) | 100 (13/13) | 75 (3/4) | 50 (1/2) |
| ↑ CRP | 12.5 (1/8) | 100 (13/13) | 100 (4/4) | 100 (2/2) |
| Abdominal symptoms | 0 (0/8) | 53.8 (7/13) | 50 (2/4) | 50 (1/2) |
| Hepatic abnormalities | 0 (0/8) | 69.2 (9/13) | 100 (4/4) | 100 (2/2) |
| Aphtosis | 0 (0/8) | 0 (0/13) | 0 (0/4) | 0 (0/2) |
| Proven gastro-intestinal inflammation | 0 (0/8) | 23.1 (3/13) | 0 (0/4) | 0 (0/2) |
| Digestive bleeding | 0 (0/8) | 30.8 (4/13) | 0 (0/4) | 0 (0/2) |
| Bleeding/ tendency to bleed | 0 (0/8) | 38.5 (5/13) | 0 (0/4) | 50 (1/2) |
| Skin rash (type) | 37.5 (3/8) | 100 (13/13) | 100 (4/4) | 100 (2/2) |
| Atopy | 37.5 (3/8) | 7.7 (1/13) | 0 (0/4) | 50 (1/2) |
| Vasculitis | 0 (0/8) | 0 (0/13) | 0 (0/4) | 50 (1/2) |
| Arthralgia/ arthritis | 0 (0/8) | 7.7 (1/13) | 75 (3/4) | 0 (0/2) |
| Central Nervous System inflammation | 0 (0/8) | 15.4 (2/13) | 25 (1/4) | 50 (1/2) |
| Other inflammatory manifestations | One case of lung infiltration with several episodes of pulmonary hemorrhages responsive to steroids | 0 | One case of bilateral pulmonary infiltrates leading to acute respiratory distress syndrome | One case of relapsing polychondritis and progressive bilateral sensorineural deafness since early adulthood |
| Clinical autoimmunity | 12.5 (1/8) | 0 (13/13) | NA | 0 (0/2) |
| Biological autoimmunity | 12.5 (1/8) | 50 (1/2) | NA | 0 (0/1) |
| Hemophagocytic lymphohistiocytosis | 0 (0/8) | 53.8 (13/13) | 25 (1/4) | 0 (0/2) |
| Hepato-splenomegaly | 25 (2/8) | 100 (13/13) | 100 (4/4) | 100 (2/2) |
| Anemia | 25 (2/8) | 100 (13/13) | 100 (4/4) | 100 (2/2) |
| Neutropenia | 37.5 (3/8) | 100 (13/13) | 25 (1/4) | 0 (0/2) |
| Thrombocytopenia | 100 (8/8) | 100 (13/13) | 50 (2/4) | 50 (1/2) |
| Abnormal platelet features | 100 (7/7) | 0 (0/2) | NA | NA |
| Abnormal bone marrow aspiration | 40 (2/5) | 91.7 (11/12) | 0 (0/1) | 100 (1/1) |
| HyperIgE | NA | 100 (3/3) | NA | NA |
| Hyper eosinophilia | NA | 100 (2/2) | NA | NA |
| Cytokinic dosage | ↑IL6, IL18, IL18BP,CXCL9 in one patient (1/1) | ↑IL6, IL1, IL18, IFNg, CXCL9 | ↑IL6, IL1, IL18, CXCL9 | ↑IL18 |
| Response to anti-IL1β | NA | Partial response for 6/8 patients<br>No response for 2/8 patients | Partial response for 3/4 patients<br>Complete response for 1/4 patients | Partial response in 1/1 patient |
| Response to anti-JAK | NA | Complete response for 1/1 patient | NA | NA |
| Alive | 87.5 (7/8) | 46.2 (6/13) | 100 (4/4) | 50 (1/2) |

NA : Not available

**FIG 1.**
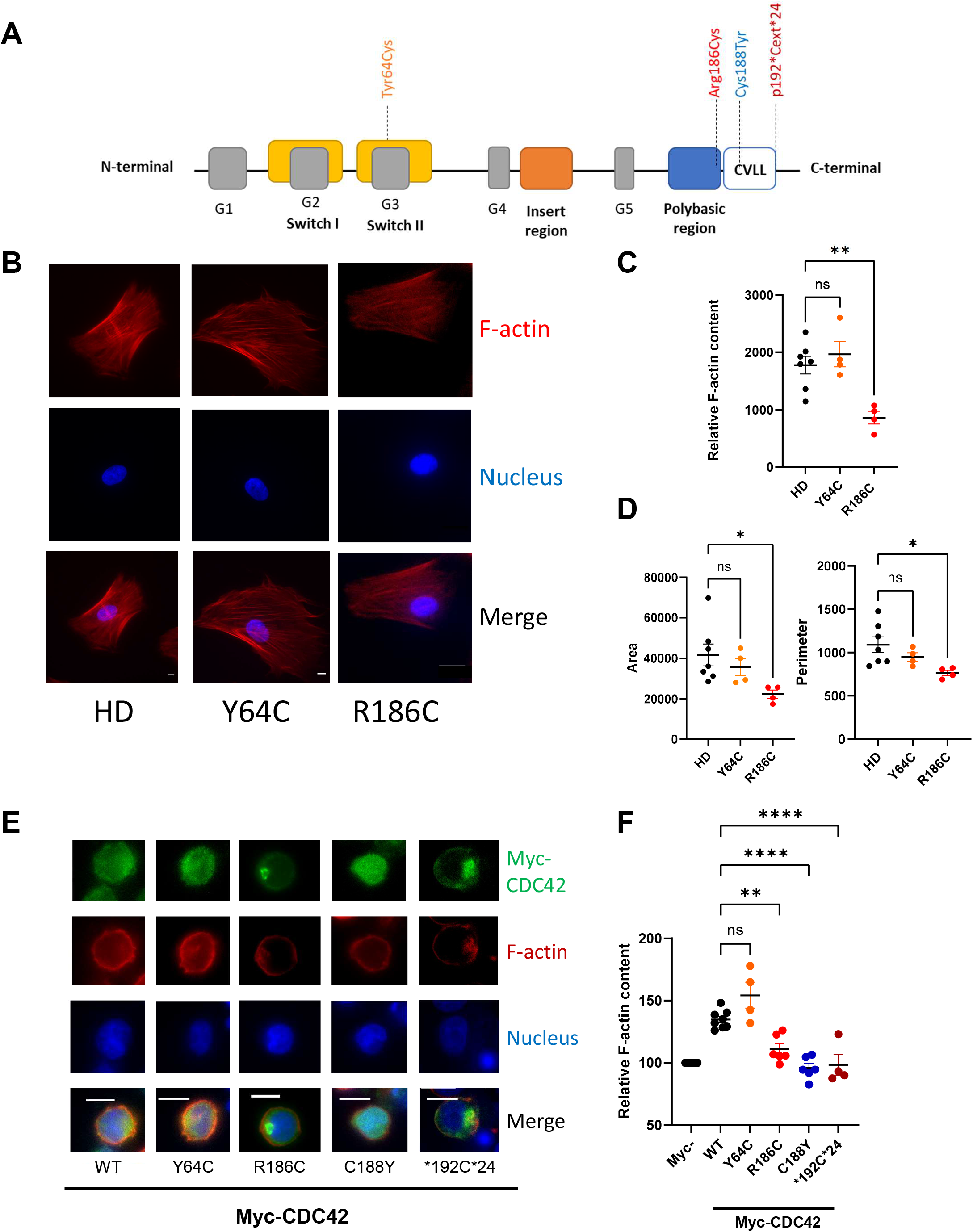
**A**, Positions of the different CDC42 mutations identified in autoinflammatory patients. **B**, Primary healthy donor (HD) or CDC42 Y64C and R186C fibroblasts labeled for actin filaments (F-actin, red) and nucleus (blue). The merged panels are shown at the bottom. Scale bars: 10 μm. **C**, Quantification of F-actin levels in patients’ fibroblasts. **D**, Quantification of fibroblasts morphological parameters: Area (left) and Perimeter (right). **E**, Subcellular localizations of myc-tagged WT, Y64C, R186C, C188Y and *192C*24 CDC42 variants in THP-1 cells. Scale bar: 10 μm. **F**, Quantification of F-actin levels in THP-1 cells expressing Myc-tagged WT or variants CDC42 based on the normalization to 100 on non-transfected Myc^-^ cells present in the cell population transfected with WT CDC42. ns: not significant.

To date, most of the molecular and signaling functional defects were only studied for the recurrent R186C variant. We and others have shown that the R186C mutation renders CDC42 defective for actin polymerization (9, 11). By contrast, the NF-κB signaling pathway was hyperactivated in CDC42 R186C expressing cells (11), in accordance with the high levels of pro-inflammatory cytokines measured in these patients (9–15). On the four CDC42 variants reported to date in autoinflammatory patients, we have studied here their impact on actin polymerization and NF-κB activation. We have also tested the possibility of a causal link between these two signaling pathways. Finally, we show that the initial classification of CDC42 autoinflammatory patients in the actinopathy group is uncorrect.

## RESULTS AND DISCUSSION

We previously reported that fibroblasts from a R186C CDC42 patient had a reduction in filamentous actin (F-actin) content (11). First, we have studied whether this result could be generalized to the Y64C TKS variant (8). For this purpose, we have stained actin filaments of primary fibroblasts from healthy donor (HD), Y64C and R186C CDC42 patients (Fig. 1B). The quantification of images showed that F-actin levels were normal in the Y64C patient whereas we could measure a decrease by half in the R186C fibroblasts (Fig. 1C). R186C CDC42 fibroblasts also exhibit a reduction in both area and perimeter compared to both HD and Y64C cells (Fig. 1D), most likely as a result of a defective spreading due to their low content in actin filaments. Thus, these results indicate that the Y64C variant identified in an autoinflammatory patient does not share the same cytoskeletal and morphological defects as the recurrent R186C variant.

Next, to test a hypothetical causal link between each of these four CDC42 variants identified in autoinflammatory patients, and actin polymerization, we ectopically expressed myc-tagged versions of them in the human monocytic THP-1 cell line (Fig. 1E). WT CDC42 was mostly diffuse throughout the cytoplasm. The C188Y CDC42 mutant tended to accumulate in the nucleus as shown before (16). Both dually lipidated R186C and *192C*24 variants accumulate in a subcellular region that was identified as the Golgi apparatus (9, 11, 14, 16). However, the N-terminal Y64C variant has a normal subcellular localization like WT CDC42. Next, WT or variants CDC42 were individually expressed in THP-1 cells and the filamentous actin (F-actin) content was measured (Fig. 1F). Results were normalized to 100 for the F-actin levels measured in THP-1 cells that do not express myc-tagged WT or mutant CDC42. Based on this quantification, we show that overexpression of WT or Y64C CDC42 induce on average a 30 to 50 % increase in actin polymerization compared to non-transfected THP-1 cells (Fig. 1F), indicating that CDC42 Y64C is fully functional. By contrast, the R186C variant was largely unable to stimulate actin polymerization, and both C188Y and *192C*24 variants were completely inactive, expressing the same level of F-actin as the non-transfected cells. Altogether, these results indicate that only the C-terminal CDC42 variants are defective for actin polymerization.

Furthermore, since we previously reported that the NF-κB pathway was hyperactivated in CDC42 R186C expressing cells (11), we next investigated the effect of the other CDC42 variants on NF-κB signaling. We first studied the nuclear translocation of NF-κB in CDC42 Y64C or R186C patients and HD fibroblasts upon LPS treatment (Fig. 2A). The majority of the NF-κB pool is cytosolic in all resting cell types. However, NF-κB nuclear translocation upon treatment with LPS was observed to a similar level in a fraction of HD and Y64C fibroblasts, and to a larger extent in R186C cells. To quantify this phenomenon, we measured the intensity of NF-κB staining in the nucleus and cytosol of randomly imaged cells (Fig. 2B). As expected, LPS stimulation elicits an increase of the nucleus/cytosol NF-κB intensity ratio for all cell types, indicating a specific nuclear translocation of NF-κB. However, although CDC42 Y64C elicits NF-κB nuclear accumulation to a similar extent as HD fibroblasts, it is clear that the R186C variant is much more efficient in driving NF-κB in the nucleus. We next studied whether NF-κB activation, that correlates with its phosphorylation, was modulated by these CDC42 variants. In the absence of LPS stimulation, the percentage of THP-1 cells that have basal phosphorylation of NF-κB was low and similar for the different conditions (below 6 %; Fig. 2C). Upon LPS stimulation, the fraction of THP-1 cells which present NF-κB phosphorylation was slightly increased in cells expressing WT or Y64C CDC42 variants. However, in the presence of C188Y, R186C or *192C*24 CDC42 variants, NF-κB activity was significantly higher compared to WT CDC42. Thus, from the four autoinflammatory CDC42 variants studied here, only the three C-terminal CDC42 variants exhibit NF-κB hyperactivation.

**FIG 2.**
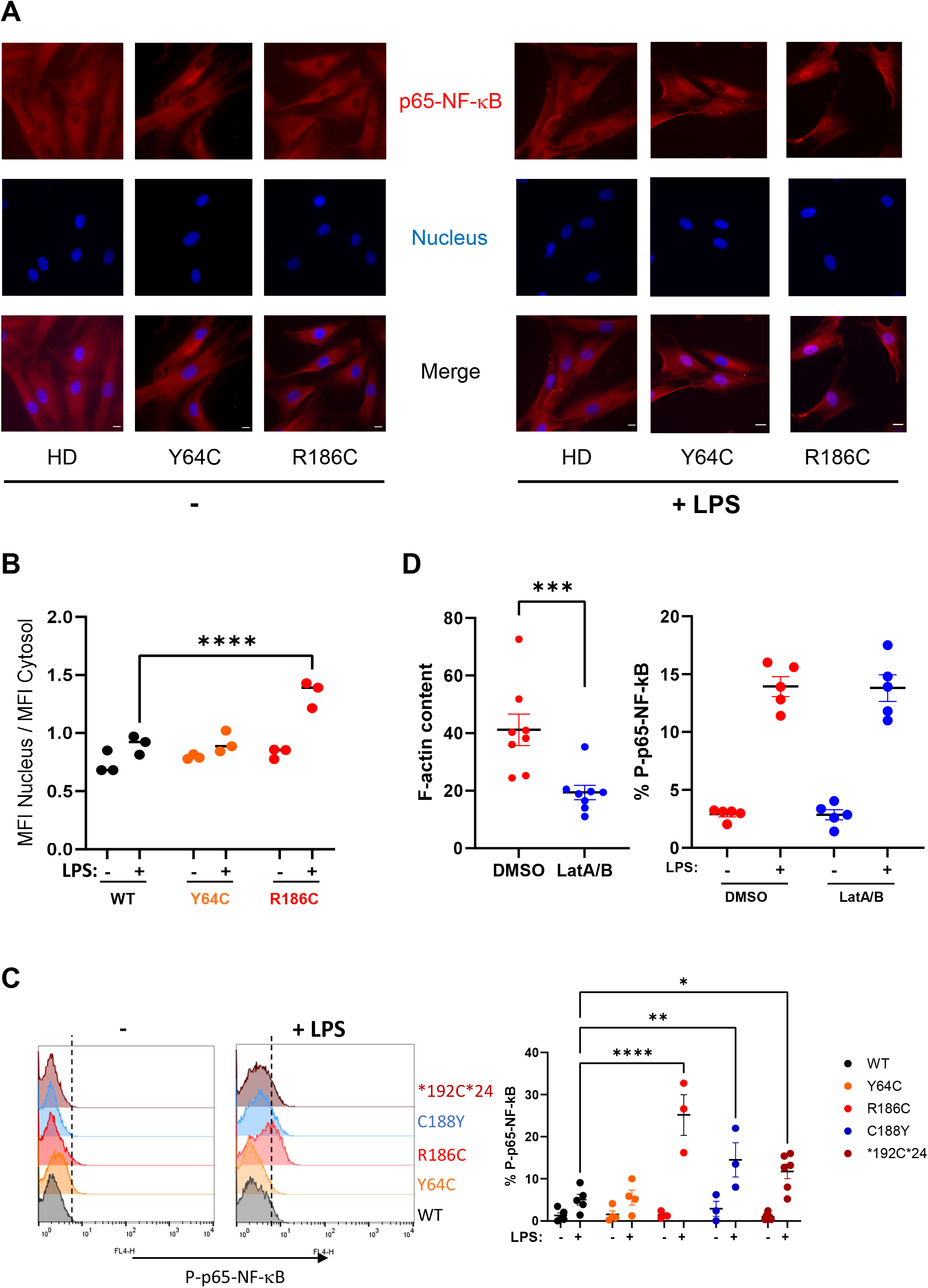
**A**, Microscopy analysis of NF-κB nuclear translocation in HD, Y64C or R186C CDC42 fibroblasts before and after stimulation with LPS. p65-NF-κB (red); nucleus (blue). NF-κB nuclear translocation appears in magenta. Scale bars: 10 μm. **B**, Quantification of NF-κB nuclear translocation measured as the ratio of nuclear to cytosolic NF-κB intensity. Each dot represents the mean value of 10 to 20 cells from one independent experiment. **C**, Left: Flow cytometry profiles of phosphorylated NF-κB in Myc-CDC42+ THP-1 cells stimulated or not with LPS. Right: Quantification of the percentage of Myc-CDC42^+^ THP-1 cells positive for Phospho-p65-NF-κB staining. Each dot represents the value from one independent experiment. **D**, Left: F-actin levels in DMSO (vehicle) or LatA/B -treated cells. Right: Percentage of Phospho-NF-κB^+^ THP-1 cells treated with DMSO or LatA/B stimulated or not with LPS.

Altogether, it is puzzling to note that all CDC42 mutants that are deficient in actin polymerization are also the ones that hyperactivate NF-κB. This inverse correlation prompted us to determine whether an intrinsic defect in actin polymerization could be responsible for triggering NF-κB activation. It has indeed been reported in the literature that pathogenic mutations in actin polymerization regulators such as WASp (18), WDR1 (19) and Hem-1 (20) can modulate inflammatory responses. Therefore, we next tested whether actin depolymerization induced by Latrunculin A or B (LatA/B) drugs could activate the NF-κB pathway. As a control, we first confirmed that these pharmacological agents were indeed efficient for decreasing the F-actin content in THP-1 cells (Fig. 2D, left). However, similar levels of NF-κB phosphorylation between the DMSO control condition and the LatA/B -treated condition were observed in both resting and LPS-stimulated cells (Fig. 2D, right). Overall, this indicates that actin depolymerization *per se* is not sufficient to induce NF-κB phosphorylation, suggesting that two divergent signaling pathways arise from CDC42 to control actin polymerization and NF-κB activation.

In this study, we show that only the C-terminal CDC42 variants are defective for actin polymerization, indicating that autoinflammatory diseases due to mutations in *CDC42* cannot be necessarily classified among the actinopathies. One common feature of these variants is their aberrant subcellular localizations (Golgi or nuclear retention) due to impairments in lipidation, as opposed to the N-terminal Y64C variant that exhibits normal distribution. Such mislocalizations could impair CDC42 binding to partners required for actin polymerization. However, we demonstrate that actin polymerization deficiency does not directly cause NF-κB hyperactivation. Again, only the C-terminal CDC42 variants hyperactivate NF-κB. This cannot be due to the sole Golgi retention of CDC42, as previously observed for Pyrin hyperactivation (14), because we show here that the nuclear C188Y variant also induces NF-κB hyperactivation. How CDC42 mislocalizations *per se* drive NF-κB hyperactivation independently of the actin cytoskeleton remains to be demonstrated, as the cross-talk between CDC42 and NF-κB needs to be clarified (21, 22). Altogether, we provide additional important clues regarding major functional defects observed in autoinflammatory patients who carry heterozygous mutations in *CDC42*. These results have obvious consequences for CDC42 patients’ management.

## METHODS

### Study oversight

Human studies were carried out according to French law on biomedical research and to the principles outlined in the 1975 Helsinki Declaration and its modifications. Institutional review board approval was obtained (DC-2023-5921 and IE-2023-3004) from the French Ministry of Higher Education and Research. All patients provided written informed consent for the conservation and use of their cells for research.

### Plasmid constructs

The plasmid pRK5-myc-CDC42 obtained from Addgene encodes for the ubiquitous isoform 1 of CDC42 and contains a myc tag in N-terminal. From this construct, CDC42 point mutants (R186C, C188Y, Y64C) were obtained by site-directed mutagenesis (Quick change kit, Agilent Technologies). The *192C*24 plasmid was produced by ThermoFisher.

### Cells

Primary fibroblasts from the Y64C patient and healthy donors were obtained from the Leuven biobank for primary immunodeficiency. Primary fibroblasts from the R186C patient and healthy donors were previously described (11). The cells were cultured in DMEM supplemented with 10 % fetal calf serum, antibiotics (penicillin and streptomycin) and sodium pyruvate. The human THP-1 monocytic cell line was provided by Serge Bénichou (*Institut Cochin*, Paris). The cells were cultured in RPMI medium supplemented with 10 % fetal calf serum, antibiotics (penicillin and streptomycin) and sodium pyruvate. All the cells were regularly tested for Mycoplasma (Lonza).

### Transfections

2×10^6^ THP-1 cells were centrifuged for 5 min at 1200 rpm and washed in 1X PBS (Gibco). They were then transfected by nucleofection with 3 μg of plasmid DNA in 100 μl of Cell Line Nucleofector Solution V (Lonza) using the V-001 program (Amaxa Biosystems). After transfection, 500 μl of complete RPMI medium warmed to 37 °C was added to the cells, which were then plated into 6-well plates, each well containing 2 mL of the same pre-incubated medium. The plates were then incubated overnight.

### Cell stimulation

LPS was purchased from Sigma (L6529) and used at 2 μg/mL.

### Intracellular staining

Cells were fixed with 4 % paraformaldehyde (Electron Microscopy Sciences) for 10 min. Then, they were washed once in the saturation buffer [PBS 1 % BSA (Sigma)] and twice in the permeabilization buffer [PBS containing 0.1 % saponin (Fluka Biochemika) and 0.2 % BSA]. Cells were then incubated for 45 min with 3 μg/ml Myc-Tag (9B11) Mouse mAb AlexaFluor 488 conjugate antibody (Cell Signaling Technology) and Phalloidin-Alexa 647 (Invitrogen, 0.5 U/ml). Nuclei were labeled with 1 μg/mL Hoechst for 10 min in the dark. Finally, the cells were washed again in the permeabilization buffer.

Quantification of the activation of the NF-κB transcription factor in THP-1 cells was performed using the same protocol as above, except for an additional post-fixation step of nuclear permeabilization of the cells with 0.1 % Triton-X100 in PBS for 10 min at room temperature, followed by two washes with the permeabilization buffer before incubation with 4 μg/ml Alexa Fluor® 647 Mouse anti-Phospho-NF-κB p65 (pS529) (BD Biosciences) for 45 min.

To evaluate the nuclear translocation of NF-κB in fibroblasts by microscopy, cells were stained with the primary anti-NF-κB p65 [(C20) SC-372; Santa Cruz Biotechnology] antibody for 45 minutes, washed twice with permeabilization buffer and incubated 45 minutes with secondary antibody. The labeling of the nuclei was next performed with 1 μg/mL of Hoechst for 5 min in the dark. Cells were finally washed twice with 1.5 mL PBS and mounted with VECTASHIELD Vibrance ® (H-1700).

### Flow cytometry

The amount of actin filaments and the percentage of active phosphorylated NF-κB in transfected THP-1 cells, were measured by flow cytometry (FACS Calibur, BD) and analyzed with FlowjoV7.

### Imaging

Images were obtained with an inverted fluorescence microscope (Eclipse Nikon TE300), a Photometrics Cascade camera, and acquired using Metamorph v7.8.9.0 software. For fibroblasts, analyses of morphological parameters, F-actin levels and NF-κB nuclear translocation were performed using Fiji software (ImageJ version 1.51u).

### Statistical analysis

Statistical analyses were performed using GraphPad Prism 10.0.1 software. Results are shown as means +/-S.E. (Standard Error) and the significance levels were calculated using the ANOVA test or the Student’s t-test. *: p<0.05, **: p<0.01, ***: p<0.001, ****: p<0.0001.

